# Retromer forms low order oligomers on supported lipid bilayers

**DOI:** 10.1101/2020.03.31.018150

**Authors:** Catherine L. Deatherage, Joerg Nikolaus, Erdem Karatekin, Christopher G. Burd

**Author notes:** Address correspondence to either author. Christopher Burd, Erdem Karatekin.

## Abstract

Retromer is a protein sorting device that orchestrates the selection and export of integral membrane proteins from the endosome via retrograde and plasma membrane recycling pathways. Long standing hypotheses regarding the Retromer sorting mechanism posit that oligomeric interactions between Retromer and associated accessory factors on the endosome membrane drives clustering of Retromer-bound integral membrane cargo prior to its packaging into a nascent transport carrier. To test this hypothesis, we examined interactions between the components of the SNX3-Retromer sorting pathway using quantitative single particle fluorescence microscopy of a reconstituted system comprising a supported bilayer, Retromer, a model cargo protein, the accessory proteins SNX3, RAB7, and the Retromer-binding segment of the WASHC2C subunit of the WASH complex. The predominant species of membrane associated Retromer are low order: monomers (∼18%), dimers (∼35%), trimers (∼24%) and tetramers (∼24%). Unexpectedly, neither cargo nor accessory factors promote Retromer oligomerization on a supported bilayer. The results indicate that Retromer has an intrinsic propensity to form low order oligomers and that neither membrane association nor accessory factors potentiate oligomerization. Hence, Retromer is a minimally concentrative sorting device adapted to bulk membrane trafficking from the endosomal system.

## Introduction

Retromer is an evolutionarily conserved protein complex that orchestrates sorting and export of integral membrane proteins from the endosome. Loss of Retromer function, which is implicated in a variety of disease conditions, results in increased rates of turnover of plasma membrane proteins and retrograde cargo proteins in the lysosome, with broad consequences to cell and organism physiology (1-3).

Retromer is composed of three proteins VPS26, VPS29, and VPS35, that form a stable, soluble heterotrimer (4-7) that is recruited to the endosome by binding to sorting signals of integral membrane protein cargo and to other membrane-associated accessory proteins, including sorting nexins, RAB7 network components, and the WASH complex (2,8). Genetic and structural analyses of Retromer trimer complexed with different sorting nexins suggests that Retromer is a modular sorting device that associates distinctly with different sorting nexins (e.g., SNX-BARs, SNX3, SNX27) to establish cargo-specific sorting and trafficking pathways (2). Despite recent insights into Retromer structure, the molecular mechanisms of Retromer-mediated sorting remain poorly understood.

We discovered that the yeast (*Saccharomyces cerevisiae*) sorting nexin, SNX3/Grd19, functions as a cargo-selective Retromer adapter that associates with Retromer on the endosome membrane and aids in cargo recognition (9). Studies of cultured human cells and other model organisms confirmed the existence of a SNX3-Retromer sorting pathway in metazoans (10-12) where cargo sorting signal is recognized via the SNX3-Retromer interface (13). Using bulk biochemical reconstitution, we previously discovered that multi-valent interactions between Retromer, SNX3, RAB7, and an integral membrane cargo confer recruitment of Retromer to the surface of small unilamellar vesicles (14). In this study we examined reconstituted components of the SNX3-Retromer sorting system on supported lipid bilayers by quantitative fluorescence microscopy. We find that when associated with a membrane, both in the presence or absence of cargo and accessory proteins, Retromer exists as monomers and lower order oligomers (dimer-to-tetramer). The results suggest that cargo is modestly concentrated by Retromer prior to export from the endosome by the SNX3-Retromer pathway, prompting a revision in long standing models of retromer-mediated cargo sorting.

## Results and Discussion

As a coat protein of endosome-derived transport carriers that recognizes retrograde cargo sorting signals, Retromer has been proposed to concentrate integral membrane protein cargo prior to transport carrier formation (1,7,15,16). To test this, we first sought to determine the oligomeric state of Retromer when it is associated with a membrane. Accordingly, a supported lipid bilayer (SLB) was constructed to mimic the relatively planar surface geometry of the vacuolar domain of the sorting endosome, where Retromer sorting domains are formed. The SLB contained physiological endosomal lipids, phosphatidylcholine (PC), phosphatidylserine (PS), and PtdIns3*P*, and non-physiological NiNTA-DGS, with a nickel ion-containing headgroup that is recognized by poly-histidine sequences, and trace amounts of rhodamine-phosphatidylethanolamine (Rh-PE) or NBD-phosphatidylethanolamine (NBD-PE), used to assess lipid mobility and the quality of the bilayer (Fig S1A). Fluorescence recovery after photobleaching (FRAP) analyses confirmed free diffusion of lipids within the supported bilayer; bilayers that were not fluid were exempt from analysis. Experiments also confirmed that association of a fluorescent His_10_-tagged peptide with the SLB is dependent upon the presence of NiNTA-DGS lipid and that the lipid-associated polypeptide is mobile (Figure S1B, C).

### Retromer exists as monomers and low order oligomers on a membrane

Retromer was assembled in lysates of bacterial cells expressing individual Retromer proteins, as we have used previously (14). To directly visualize Retromer on the SLB by fluorescence microscopy, VPS26 was produced with N-terminal His_10_ and SNAP tags (Figure 1), which facilitated binding to the SLB and labeling with a fluorescent dye (AlexaFluor488), respectively. Incorporation of His_10_-SNAP-VPS26 fusion protein into the Retromer trimer was indistinguishable to that of VPS26, indicating that Retromer structure is maintained in the tagged protein. After incubating labeled Retromer with the SLB for 2 hours, the SLB was washed to remove unbound proteins and total internal reflection fluorescence microscopy (TIRFM) was used to visualize SLB-associated proteins (17). The results show that Retromer is distributed homogenously on the SLB (Fig. S1C), suggesting that it does not self-organize into clusters over a range of concentrations (1 nM nominal Retromer concentration is shown in the figure).

**Figure 1.**
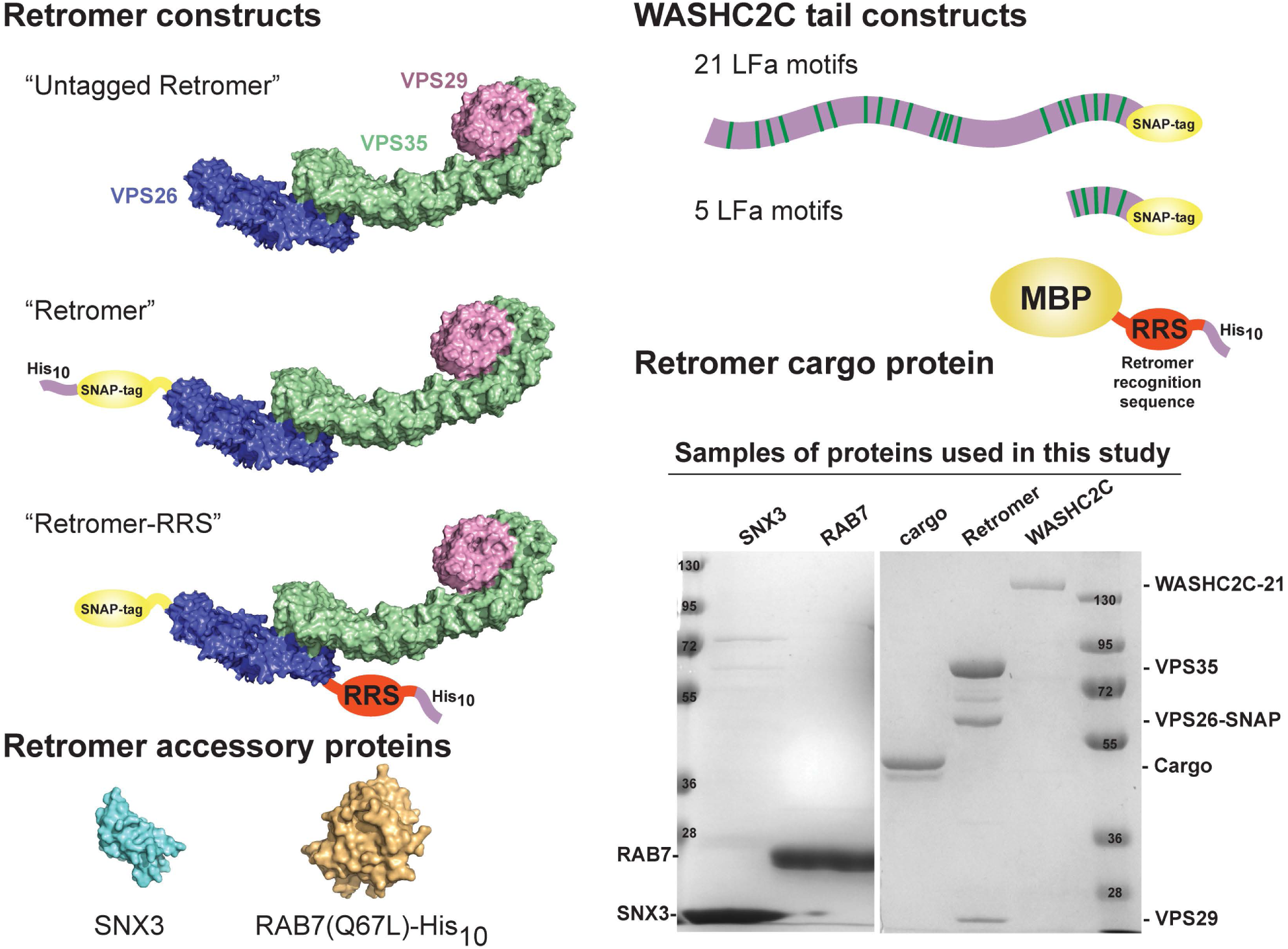
Protein constructs used in this study. “Untagged Retromer” refers to recombinant VPS26/VPS35/VPS29 Retromer trimer. “Retromer” refers to a trimer where VPS26 carries a SNAP-tag (for labeling with AF488 dye) and a His_10_ tag for attachment to supported bilayers containing NiNTA-DGS lipids. “Retromer-RRS” contains a VPS26 subunit with a SNAP tag at the N-terminus and the sequence of the DMT1-II Retromer sorting signal (“RRS”) and a His_10_ tag fused at its C-terminus. In this construct, stoichiometric Retromer-cargo interactions are enforced artificially. SNX3, RAB7(Q67L)-His_10_ (a GTP-locked, constitutively active mutant), WASHC2C constructs contain either 21 or 5 LFa motifs. “Retromer cargo protein” is composed of an N-terminal maltose binding protein with a sorting signal and a His_10_ tag fused at its C-terminus. An example of a Coomassie blue-stained gel shows the purity and stoichiometry of complexes. The following structure files from the Protein Data Bank were used to prepare the figure: RAB7A, 1T91; SNX3, 2YPS; Retromer, 6H7W.

Next we established a single molecule fluorescence microscopy assay to determine Retromer oligomeric state on the SLB. These experiments use lower protein concentrations in the medium (∼75 pM), resulting in a lower protein density on the SLB, such that a minimum of 3-4 pixels (∼0.75-1 micron) separation between fluorescence labeled particles is favored, facilitating single-particle analysis. We monitored intensity of AF488-labeled Retromer puncta continuously over time at high frame rate (17-18 ms exposure per frame) to capture single fluorophore bleaching steps until most puncta bleached away (Fig. 2A). Fluorescent particles photobleached in either single or multi-step bleaching profiles, indicative of either single or multiple fluorophores in the particle (Fig. 2B). We used the decrease in fluorescence intensity due to the last bleaching event in individual puncta to estimate the intensity of single AF488 fluorophores (Fig. 2B) and then to calculate Retromer copy number per particle. Importantly, in this procedure every single image stack provides its own single-molecule intensity calibration for the cluster copy number estimation, enhancing robustness of the approach against experiment-to-experiment variations.

**Figure 2.**
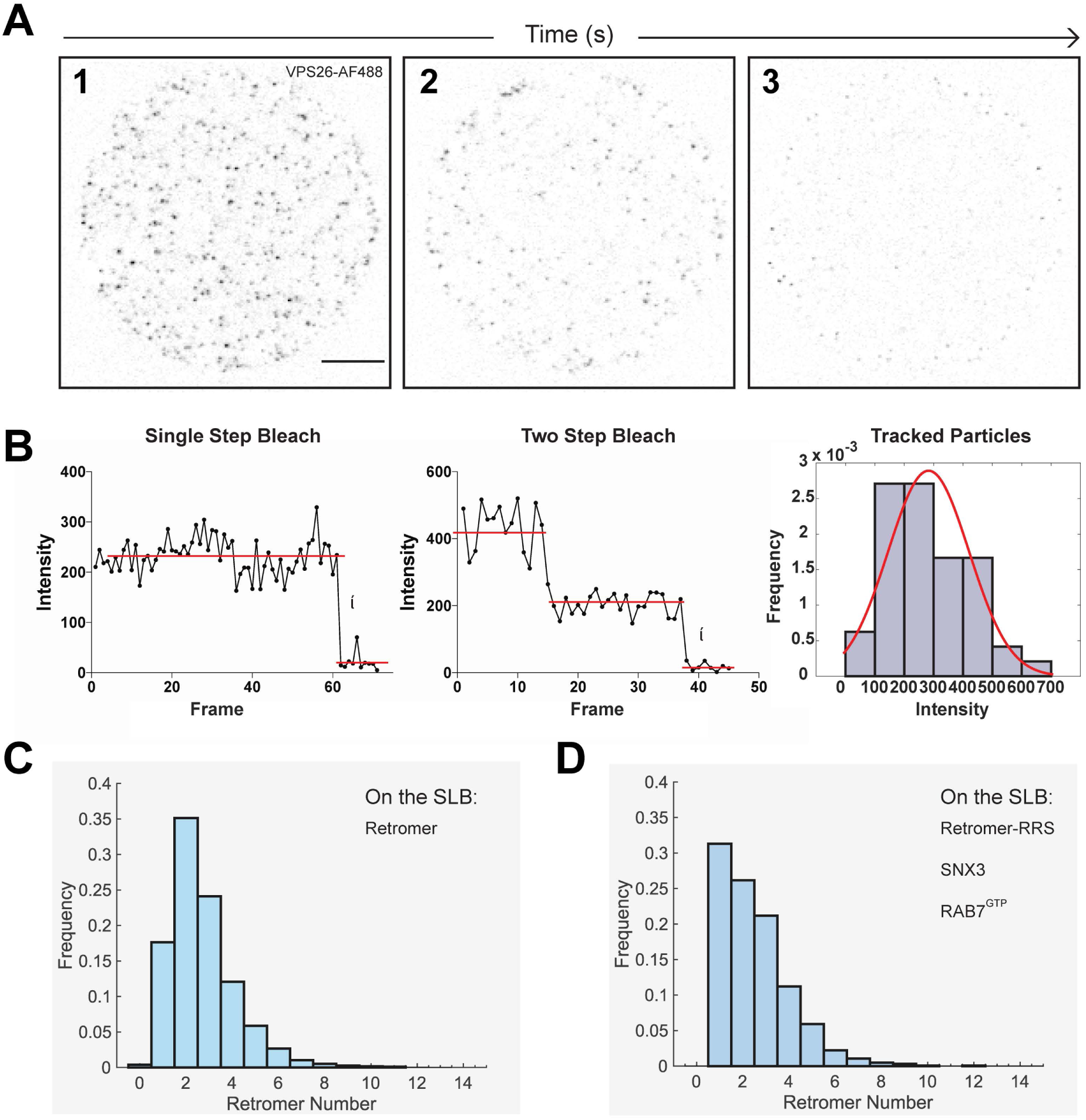
Single-particle analysis of Retromer clusters on supported bilayers. **(A)** Shown are frames from a TIRFM image stack of AF488-labeled Retromer (solution concentration 75 pM; gray values inverted). After focus was established at another region of the SLB, auto-focus was turned on. Acquisition started after moving to a new, unbleached SLB region in dark. Thus, the spots in the first frame (image 1) represent intact Retromer clusters that have not yet started bleaching. The movie was acquired at a high frame rate (∼17.8 ms per frame) such that single-molecule fluorescence bleaching events could be captured. Fluorescent spots were tracked using the SpeckleTrackerJ plugin of ImageJ, usually starting from a frame after some of the spots bleached away (image 2), which facilitated tracking. Imaging continued until nearly all spots disappeared (image 3). A blurred version of the last frame was subtracted from all previous frames to correct for uneven illumination and background during analysis. Scale bar: 10μm. **(B)** Examples of fluorescence intensity profiles of single-particle tracks and distribution of single-fluorophore intensities. Left: the mean pixel value from a 3×3 pixel region centered around the tracked position of a spot as a function of time (frames). Because it was easier to identify the final bleaching step, we only used the final drop in intensity to estimate single-molecule intensities. A distribution of fluorescence bleaching events are shown on the right with a Gaussian distribution overlaid (red line). **(C)** Frequency distribution of the number of Retromer complexes per cluster on a SLB. The first frame from movies as in A) were used to calculate intensities of fluorescent spots. These intensities were converted to the number of Retromer per cluster, using the single-fluorophore intensity calibration (obtained as in B), and the labeling efficiency. **(D)** His-tagged, AF488-labeled Retromer-RRS (solution concentration 75pM), SNX3 (3.25nM), and RAB7(Q67L)-His_10_ (1nM) were attached to the SLB and analyzed by single-particle fluorescence. Although there are differences in the distribution of Retromers per cluster compared to Retromer alone (C) (two-sample Kolmogorov-Smirnov test, p<0.01), neither the mean nor the span of the two distributions indicate a biologically meaningful shift to higher oligomeric states (average of 2.5 vs 2.7 Retromers per cluster, span=0-13 vs 0-12, for C and D, respectively).

Results for independent preparations of retromer were pooled together. The combined results of four independent Retromer preparations are plotted as a histogram showing the frequency distribution of Retromer complexes on the SLB (Fig. 2C). The predominant species of Retromer on the SLB are monomers (∼18%), dimers (∼35%), trimers (∼24%) and tetramers (∼13%). Also observed are pentamers and rare higher-order oligomers up to ten (∼10%). This distribution of Retromer oligomers on SLBs is similar to that reported in a cryo-electron microscopy study where dimers and tetramers were also the prevalent Retromer species in vitrified ice (18). These results suggest that membrane association *per se* does not influence Retromer oligomeric sate.

### Retromer accessory proteins do not influence oligomeric state

Retromer association with the endosome membrane is conferred by binding to integral membrane cargo proteins, sorting nexins, and RAB7 (14,19-21), therefore we next sought to determine if these accessory factors influence Retromer oligomerization on a SLB. His-tagged, AF488-labeled cargo protein was homogeneously dispersed on the SLB (Fig. S2A) and incubation with “untagged Retromer” (recombinant, not his-tagged or labeled) did not affect its appearance (not shown). However, the effect of Retromer may not be apparent due to the low affinity with which Retromer binds sorting signals. Accordingly, we added the sorting nexin, SNX3, which facilitates Retromer membrane recruitment and forms part of the DMT1-II cargo binding site (13), onto the SLB. After confirming that purified, labeled SNX3 associates with the SLB by binding to PtdIns3*P* (Fig. S1D), SNX3 and untagged Retromer were sequentially added in stochiometric excess of the nominal cargo concentration and the distribution of cargo fluorescence was monitored for any changes occurring with the addition of SNX3 and soluble Retromer (Fig. S2). At all times examined, cargo fluorescence was homogenously distributed on the surface of the SLB (Fig. S2B). Thus, at the protein concentrations accessible in our experimental system, a model Retromer cargo is not clustered by SNX3-Retromer (Fig. S2B, C).

We next measured Retromer particle size at low protein density on the SLB using quantitative single-particle TIRFM. For these experiments we used a modified system because at the low densities of proteins on the SLB required for single-particle analysis, only a small proportion of cargo molecules will be bound by Retromer. Accordingly, we constructed a non-dissociable model of the Retromer-cargo complex, termed “Retromer-RRS”, that was inspired by crystallographic studies of Lucas et al. (13) where a similar VPS26-DMT1-II fusion protein facilitated elucidation of the cargo binding site on SNX3-Retromer (13). The effect of an additional Retromer accessory protein, RAB7, was also examined because RAB7 is proposed to aid in Retromer recruitment to the endosome (14,20). His-tagged, AF488-labeled (via the SNAP tag) Retromer-RRS fusion protein, SNX3, and RAB7(Q67L)-His_10_ (a mutant form of RAB7 in the GTP-bound conformation) were incubated together with the SLB. Following washing of the SLB to remove unbound protein, single-particle fluorescence data were collected to determine the distribution of Retromer-RRS complexes present on the SLB (Fig. 2D) as before. In the presence of SNX3 and RAB7 the mean oligomeric state of Retromer-RRS is 2.5 (Fig. 2D). This modest decrease in mean cluster size (2.7 vs. 2.5 Retromer complexes) indicates that Retromer oligomerization on the SLB does not require, and is not significantly influenced by, the presence of SNX3, RAB7, or by occupancy of its cargo-binding site. Our conclusion contrasts with that of a study of yeast Retromer and accessory proteins interacting on the surface of a giant unilamellar vesicle (GUV) which concluded that ySNX3 and cargo potentiate clustering of Retromer and associated proteins on the GUV membrane (19). This difference might be attributed to the different membrane models used (rigid SLB vs GUV), however, we note that non-quantitative methods were used to infer changes in relative protein abundances across the GUV membrane, and that clustering of yeast SNX3-Retromer depended on the presence of a second, non-physiological membrane binding site (a His_6_ tag) on ySNX3 (19).

### WASHC2C disordered segment does not influence Retromer oligomeric state

In metazoans, Retromer recruits the WASH protein complex from the cytosol to the endosome membrane where it promotes ARP2/3-dependent actin polymerization and Retromer-dependent sorting (8,22-25). Binding of WASH to Retromer is conferred by an approximately 1100 amino acid long unstructured segment of the WASHC2C/FAM21C subunit containing 21 “Leucine-Phenylalanine-acidic motifs (“LFa”: L-F-[D/E]_3-10_-L-F) shown to constitute Retromer-binding sites by solution-phase binding assays (24,26). On the basis of the multi-valency of WASHC2C-Retromer interaction, Jia et al. (24) speculated that recruitment of WASH to the endosome membrane would result in clustering of membrane associated Retromer-cargo complexes (24). We tested this hypothesis using our single particle analysis platform.

We first determined if SLB-associated Retromer can recruit WASHC2C-21 (Fig. 1), composed of an ∼1100 amino acid long segment of WASHC2C containing all 21 LFa motifs and a C-terminal SNAP tag, to the SLB (Fig. 3). Labeled WASHC2C-21 was incubated alone, or co-incubated with labeled His_10_-tagged Retromer, with a SLB. We confirmed that WASHC2C-21 binds to the SLB via Retromer by imaging both proteins on an area of the SLB with enriched retromer using multiple fluorescence channels (Fig. 3). We then asked if segments of the WASHC2C unstructured segment can influence Retromer cluster size on the SLB. His-tagged, AF488-labeled Retromer was attached to the SLB via binding to NiNTA-DGS and then incubated with varying amounts of WASHC2C-21 to cover stoichiometries ranging from Retromer excess to WASHC2C excess (50:1, 20:1, 5:1, 1:1, and 1:13) (Fig. 4, WASHC2C-21)). A truncated version, WASHC2C-5, containing just the last five LFa motifs was also examined (Fig. 4, WASHC2C-5). The results show that the number of Retromer complexes in single particles did not vary substantially over a broad range Retromer:WASH ratios for both WASHC2C-5 and WASHC2C-21. Curiously, we saw with different Retromer:WASHC2C-5 ratios (50:1 and 1:13) the proportion of Retromer monomer was increased, possibly indicating that this fragment of WASHC2C interferes with Retromer oligomerization or complex stability. These results indicate that, under the conditions tested, the WASHC2C unstructured segment does not elicit clustering of Retromer.

**Figure 3.**
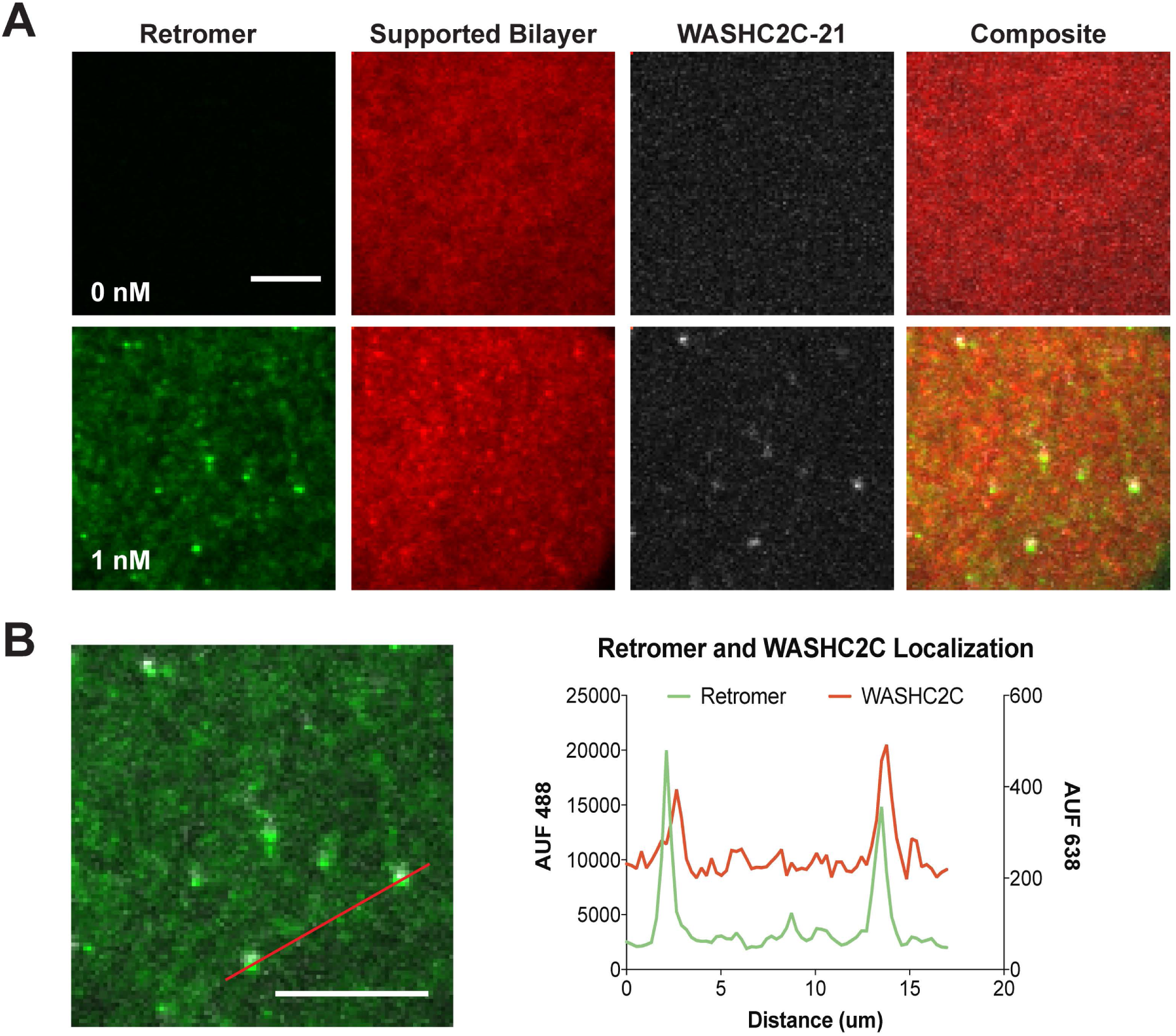
WASHC2C is recruited to the supported bilayer by Retromer. **(A)** SLBs were incubated with AF647-labeled WASHC2C-21 (solution concentration 1nM) and no Retromer (top row), or with AF488-labeled Retromer (solution concentration 1nM) (bottom row). After washing, Retromer, the SLB (RhodaminePE), and WASHC2C were imaged by TIRFM; SLBs were incubated with the WASHC2C C-terminal tail domain (AF647-labeled WASHC2C-21) (1nM). WASHC2C-21 fluorescence is seen only on SLBs with attached Retromer. **(B)** Image and line scan of Retromer and WASHC2C TIRFM channels on the SLB. Scale bar: 10μm.

**Figure 4.**
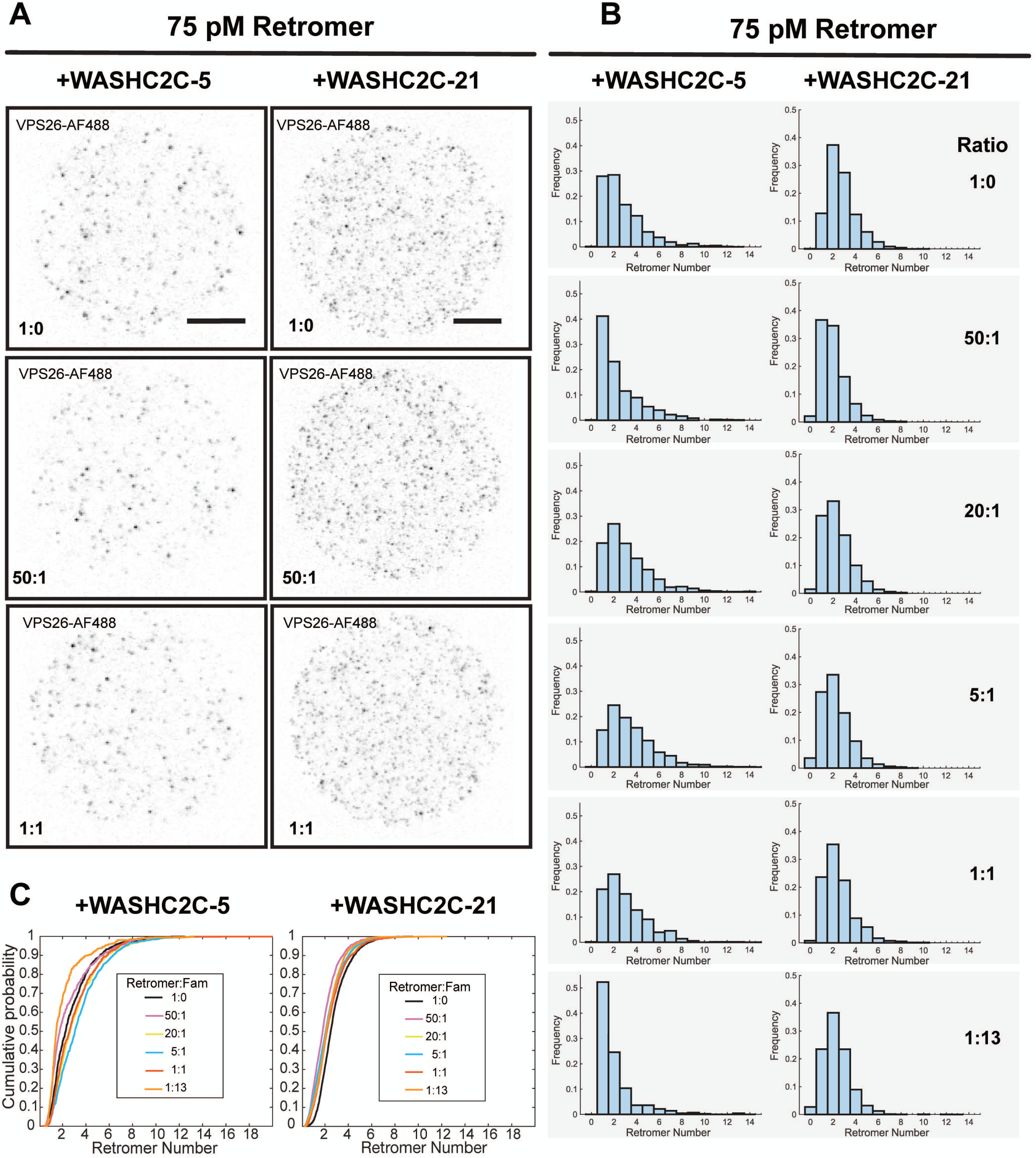
WASHC2C does not influence Retromer oligomeric state. His-tagged, AF488-labeled Retromer was attached to a SLB (solution concentration 75pM) and AF647-labeled WASHC2C protein was then added to the indicated Retromer:WASHC2C ratio. **(A)** Retromer fluorescence is shown (gray values inverted). The left column shows incubations with WASHC2C-5 (containing 5 LFa motifs) and the right column shows incubations with WASHC2C-21 (containing 21 LFa motifs). Representative micrographs of a subset of reaction conditions are shown. **(B)** Distributions of Retromer puncta incubated with WASHC2C-21 at the indicated ratios. **(C)** Cumulative distribution function (CDF) plots for each Retromer-WASHC2C condition tested.

Finally, we asked if Retromer, cargo, SNX3, RAB7, and WASHC2C act synergistically to influence Retromer clustering at low protein densities (Fig. 5). As before, Retromer-RRS was used to enforce cargo occupancy. In the absence of any other factors, the distribution of Retromer-RRS oligomers (in the presence of SNX3 and RAB7^GTP^) is similar to the distribution of Retromer oligomers (Fig. 2). Next, WASHC2C-21 was added to the reactions to examine the effect of sub-stoichiometric (50:1) and stoichiometric (1:1) WASHC2C-21. Single particle analyses revealed small increases in the proportions of Retromer-RRS monomer at both Retromer-RRS:WASHC2C-21 ratios, similar to the effect of sub-stoichiometric amounts of WASHC2C-5 (Fig. 4). These results indicate that binding of Retromer and WASHC2C on the SLB is (or is close to) stoichiometric, and further suggest that WASHC2C does not exert an effect (e.g., allosteric) to promote Retromer oligomerization. Consistent with these interpretations, Jia and colleagues reported that only the last two of the 21 LFa motifs (LFa_20-21_), which are those bound by VPS35 with the highest affinity, are essential for WASH-dependent sorting of integral membrane cargo (24). These findings suggest that WASH does not exert its role in the Retromer pathway by clustering Retromer or Retromer-cargo complexes.

**Figure 5.**
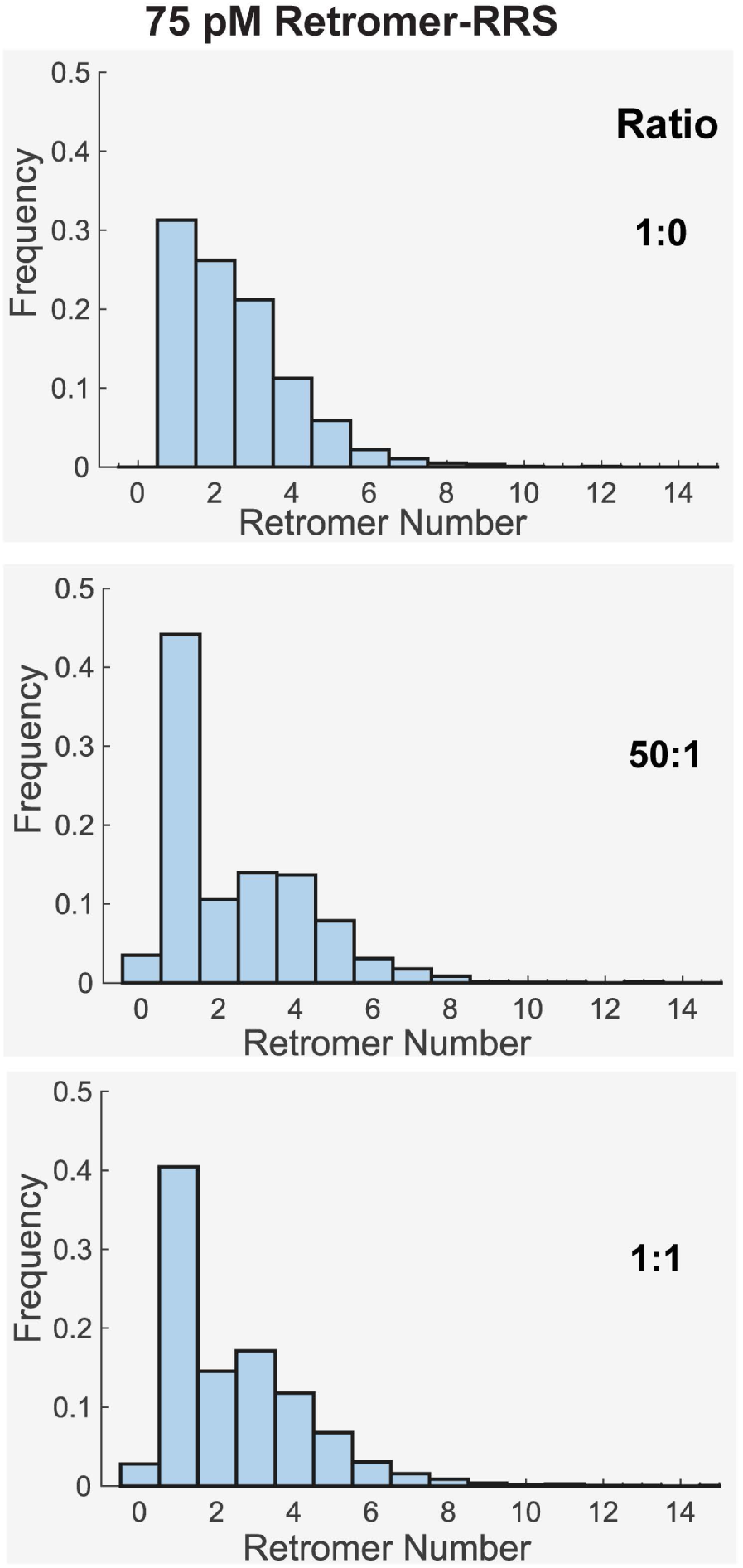
Cargo, SNX3, RAB7 and WASHC2C do not influence Retromer oligomeric state. Retromer-RRS (AF488-labeled; solution concentration 75pM), SNX3 (1nM) and Rab7(Q67L)-His_10_ (1nM) were incubated with a SLB. Following washing to remove unbound protein, WASHC2C-21 was added to the reaction cell at the indicated Retromer:WASHC2C-21 ratio. TIRFM analysis of single Retromer particles was used to measure the number of Retromer complexes in individual puncta. Note that the data in the top panel is also presented in Figure 2D.

### Implications for Retromer sorting mechanism

When associated with a membrane, human Retromer has an intrinsic propensity to form low order clusters (<5 monomers) and neither membrane association, nor the membrane-associated accessory factors - cargo, SNX3, RAB7, and WASHC2C - potentiates Retromer oligomerization on a SLB. The sizes of Retromer clusters observed in this study agree well with conclusions of biophysical and structural studies of Retromer alone in solution-phase, where Retromer monomer and dimer were the most prevalent species (13,18,27). In our study, Retromer dimers were generally the most abundant species observed on SLBs and this likely reflects two different Retromer-to-Retromer binding modes, where one is the 2-fold symmetric dimer observed in solution phase described by Kendall and colleagues (18), and the second mode is mediated by the VPS35-VPS35 dimerization interface observed in the Retromer-SNX-BAR coat (28) and in solution (13). A distinct binding mode(s) is needed to explain the small proportion of higher order oligomers observed in our study, which might consist of chains Retromer complexes in solution-phase observed by cryo-electron microscopy (18).

Retromer has been proposed to constitute a coat protein complex for endosome-derived transport carriers that, by analogy to better characterized conventional vesicle coats (e.g., clathrin) that polymerize on the membrane to enrich nascent carriers in particular integral membrane cargo (7,16). A key feature of the conventional paradigm is the small area of coated membrane that gives rise to small (<100 nm diameter) transport carriers. In contrast, sorting of retrograde and recycling integral membrane in the endosomal system is weakly concentrative, relying instead on the large surface area of endosome-derived carriers to mediate bulk export of proteins and lipid from the endosome (29). Results presented here suggest that Retromer concentrates cargo minimally, though this study was necessarily limited so it is possible that conditions or factors not examined, such as different membrane topologies, rigidities, and/or WASH-mediated actin polymerization, influence Retromer oligomerization. We note, however, that the structure of yeast VPS5-Retromer coated membrane tubules that show the coat to be highly heterogenous with limited long range order, which is consistent with a minimally concentrative sorting mechanism (2,28).

Collectively, data do not support longstanding hypotheses of Retromer sorting that invoke oligomerization as a driving force for sorting of integral membrane proteins in the endosomal system. Rather, low order oligomerization of Retromer and associated factors is likely an adaptation of bulk membrane trafficking pathways characteristic of the endosomal system that ensure membrane homeostasis of the plasma membrane and endo-lysosome organelles.

## Materials and Methods

### Recombinant Proteins

cDNAs encoding human SNX3, RAB7, VPS35, VPS26, VPS29, VPS26-RRS, WASHC2C-21 (WASHC2C-357-1318) and WASHC2C-5 (WASHC2C-921-1318), and model cargo protein were amplified by PCR and cloned into bacterial expression vectors. Retromer, RAB7, and SNX3 were prepared as described in Harrison et al. (14). Fluorescence-tagged Retromers, Retromer-RRS, model cargo protein, and WASHC2C fragments were prepared as described here.

Proteins were prepared fresh on the first or second day of each three-day experiment. Proteins were expressed by auto-induction (30) from BL21(DE3) *E. Coli* cells. The cells were pelleted from culture medium and re-suspended in lysis buffer. Cells were lysed by three passages through a cell disruptor (Avestin) at >10,000 psi. The lysate was clarified by 30,000xg centrifugation. Lysis buffer #1 [20 mM 4-(2-hydroxyethyl)-1-piperazineethanesulfonic acid (HEPES) at pH 8, and 150 mM NaCl] was used for Retromer, Retromer-RRS, and SNX3. Lysis buffer #2 (with higher salt and glycerol) [25 mM HEPES, 500 mM NaCl, 20 mM imidazole, 10% glycerol, 1 mM TCEP, pH 7.8] was used for model cargo protein, the WASHC2C fragments, and RAB7. Both lysis buffers were supplemented with 0.1 mM 4-(2-Aminoethyl) benzenesulfonyl fluoride hydrochloride, 1 mM DTT, and cOmplete™ protease inhibitor cocktail tablet (Roche Diagnostics).

GST-tagged proteins (Retromer, SNX3, WASHC2C-5 and WASHC2C-21) were purified by incubating clarified lysate with glutathione (GSH) Sepharose 4B (GE Healthcare) beads. The beads were washed with lysis buffer and the proteins released from the beads with TEV protease. If the protein was to be fluorescently labeled, the labeling was carried out before the proteins are released from the beads. Purified proteins were quantified by Bradford assay (Thermo Scientific).

Retromer: Retromer subunit VPS35 was GST-TEV-tagged. The other Retromer subunits (untagged VPS-29, untagged VPS26, his-SNAP-tagged VPS26, and his-SNAP-tagged VPS26-RRS fusion) were not GST-tagged. Cell pellets of separate cultures expressing VPS35, VPS29, and one of the VPS26 constructs were combined and resuspended together in lysis buffer #1. After cell lysis and clarification, Retromer was purified using the GST-tagged protein protocol, where the VPS35-conjugated beads were washed with lysis buffer #1. Unpartnered VPS26 and VPS29 subunits were washed away during purification. Normal assembly of heterotrimeric Retromer complexes in lysis buffer was confirmed by SDS-PAGE.

SNX3: GST-TEV-SNX3 was purified as described for Retromer, using lysis buffer #1. WASHC2C-21. GST-TEV-WASHC2C-357-SNAP was purified as described for Retromer but using lysis buffer #2.

WASHC2C-5. His_6_-SUMO-WASHC2C-5-SNAP was purified using lysis buffer #2 and Ni-NTA-agarose beads (Qiagen). The protein was incubated with Ni-NTA beads and washed with 50 mM imidazole. The His_6_-SUMO-WASHC2C-5-SNAP was released from the Qiagen beads with His6-SUMO protease.

RAB7: His_10_-RAB7(Q67L)-His_10_, a constitutively active form of the protein (31), was purified using an ÄKTAprime plus fast protein liquid chromatography (FPLC) system equipped with a 1-mL His-Trap HP column (GE Healthcare-Amersham Biosciences) and the high salt buffer. Immediately before use, RAB7 was incubated on ice with a 5X excess of 100mM GTP disodium salt solution (Sigma) for 30 minutes. In all experiments with RAB7-GTP, the RAB7-GTP is His-tagged and attaches to SLBs by binding to nickel lipids.

Cargo: Model cargo protein was constructed from the cytoplasmic tail of the Retromer cargo (amino acid 532-568), DMT1-II (12,14) with a His_10_ tag followed by an 18 amino acid linker containing a cysteine (used for fluorescence labeling) (32), a 37 amino acid fragment of the divalent metal transporter DMTII-1 containing a Retromer recognition sequence (12,14), followed by maltose binding protein (MBP). The model cargo protein was purified by the same procedure as RAB7.

### Fluorescence labeling of recombinant proteins

Proteins were labeled on the second day of a three-day experiment. SNAP-tagged proteins were fluorescence labeled while still bound to purification beads by incubation with a 2X molar excess of SNAP Surface Alexa Fluor 488 or 647 as marked overnight at 4°C with constant mixing. After 16-18 hours, the unreacted dyes were removed by thoroughly washing the beads with a minimum of 60 bead-volumes of lysis buffer prior to protein release. After free dye removal, all proteins clarified with a 100,000x*g* centrifugation step and immediately quantified by Bradford assay. The concentration of dye was determined with a spectrophotometer. The labeling efficiency was calculated by *e* = *M*_*dye*_*/M*_*protein*_, where *M*_*dye*_ and *M*_*protein*_ are the molar concentration of dye and protein. Proteins with labeling efficiency below 75% were not used.

Model cargo protein was labeled via the engineered single cysteine in the linker between the His-tag and the cargo protein sequence. The protein was labeled with a 5X excess of AlexaFluor488 C5 maleimide dye (ThermoScientific) in the dark and tumbled overnight at 4°C. Unreacted dye was quenched with the addition of 5 mM 2-Mercaptoethanol (Sigma-Aldrich). Free dye was removed with 3 sequential 2 ml Zeba desalting columns (ThermoScientific) that had been equilibrated with liposome Buffer A [25 mM HEPES, 200 mM NaCl, 1mM MgCl_2_, pH 7.5] with 10% glycerol. To label SNX3, a single cysteine (L11C, C140S) mutant of GST-SNX3 was labeled overnight on the GSH sepharose beads (GE Lifesciences) at 4°C with a 5X excess of AlexaFluor546 C5 maleimide dye (ThermoScientific). Unreacted dye was quenched and removed as described.

### Liposomes

Liposomes were prepared generally by the protocol of Su et al. (2017) (33) with some adjustments. Liposomes were made from pure synthetic lipids (Avanti Polar Lipids, and Echelon Biosciences (PtdIns3*P*)) by combining 92 mol% dioleoyl-phosphatidylcholine, 5 mol% dioleoyl-phosphatidylserine, 2 mol% [1,2-dioleoyl-sn-glycero-3- [[N- (5-amino-1-carboxypentyl) iminodiacetic acid]] succinyl (nickel salt) (NiNTA-DGS)], and 1 mol% Di-palmitoyl Phosphatidylinositol 3-phosphate in a new 4 ml glass vial. Trace amounts (0.01*μ*l) of either 1,2-dipalmitoyl-sn-glycero-3-phosphoethanolamine-N-(lissamine rhodamine B sulfonyl) (ammonium salt) (Rh-PE) or 1,2-dipalmitoyl-sn-glycero-3-phosphoethanolamine-N-(7-nitro-2-1,3-benzoxadiazol-4-yl) (ammonium salt) (NBD-PE) (Avanti) were added to visualize the SLB. For a typical total volume of ∼80 *μ*l of lipids in the vial, one ml of 9:1 chloroform:methanol was added to the vial and the contents were gently swirled to mix the different lipids together. The contents of the vial were dried to a lipid film with nitrogen and residual solvent removed for 2 hours under vacuum. The lipid film was rehydrated in the vial to 0.62 mM with 400*μ*l of Buffer A by shaking for 30 minutes and was then transferred to an Eppendorf tube. The lipid solution was then frozen at −80°C until the day of the experiment. On the day of imaging, lipid-buffer mix was sonicated on ice with a microtip sonicator (Misonex, Qsonica (S-4000, 417A)) for 20 minutes at an amplitude of 30 and 50% duty cycle. The sonicated liposomes were centrifuged for 20 minutes at 100,000xg to pellet any larger liposomes or aggregates. The supernatant was used to make the SLB within 30 minutes of clarification.

### Supported lipid bilayers

SLBs were made by the protocol of Su et al. (2017) (33) with some adjustments. A 96-well, black, glass-bottomed plate that has low background fluorescence (Matriplate MGB096-1-2LG-L) was cleaned in an overnight soak in 5% Hellmanex III heated to 50°C. After thoroughly rinsing with Mili-Q-filtered water and drying the slide, the clean wells are sealed with PCR sealing foil sheets (ThermoScientific). To form an SLB, an individual well was opened and cleaned with two 1-hour incubations at 50°C of freshly prepared, sterile filtered 5M NaOH. The well was rinsed twice with 0.5 ml Mili-Q water and twice with Buffer A. The rinsed well was then filled with 0.2 ml of Buffer A and 10 *μ*l of the liposomes were added. After incubation for one hour at 37°C, the well was washed three times with buffer A to remove any non-adhered liposomes. The SLB was then blocked with 0.1% casein in Buffer A for 20 minutes at 37°C. We checked the mobility of the SLB by FRAP or visual inspection (described below) for each experiment; immobile membranes were discarded.

In one three channel co-localization experiment (Fig. 3), the fluorescence signal was too weak for visualization on the SLB alone because of the low affinity between Retromer and WASHC2C. In this experiment, we looked at the bright spots where more lipid and thus protein was concentrated, and signal-to-noise was high enough to detect co-localization.

### Protein attachment to supported bilayers

His_10_-tagged proteins were bound to NiNTA-DGS lipids in the SLB. Proteins were added to the well containing 0.2ml of Buffer A with 0.1% casein and 1 mM TCEP. The protein and the bilayer were incubated in the well in the dark for two hours at 30°C to get a secure attachment to the bilayer. After protein addition, the bilayer was washed 3X with 0.1% casein Buffer A. Proteins not binding through His_10_ tags (SNX3, Retromer, and WASHC2C), were incubated with the SLB for 30 minutes in the dark at room temperature and then washed 3X with 0.1% casein Buffer A. Wells with immobile proteins were not used.

### Fluorescence microscopy

TIRF images were collected on a custom-build polarized TIRF microscope with an Olympus microscope body, a 60x objective (oil, PlanApo, NA 1.45), an Andor EM CCD camera (Ixon Ultra), and Micro-Manager software (34). Lasers at 488 nm, 561 nm, and 638 nm were used to excite NBD-PE/Alexa Fluor 488, Rh-PE/Alexa Fluor 546, and Alexa Fluor 647, respectively. Single micrographs were collected with a 100 ms exposure time and movies were imaged with a 17.74 ms frame duration. The 488 nm laser for FRAP measurements with NBD-PE was used at maximum laser power and data analyzed as described in (35,36).

### Readouts of large-scale clustering events

Some systems, for example the reconstituted adhesion receptor Nephrin signaling system (32), may undergo large-scale phase transitions in which very large clusters, networks, or coats of proteins may form. To identify clustering events larger than ∼10 particles we prepared SLBs with high protein densities and examined them by TIRF fluorescence microscopy. These experiments used high protein concentrations in the medium (∼1nM), resulting in high protein densities on the SLB, with < 1 pixel (∼0.25 *μ*m) separation between fluorescence labeled proteins. Images were inspected visually for localized puncta of intense fluorescence. In some more sensitive quantitative experiments, intensity measurements (line scans) across the illuminated field of view were used to assess fine-scale variation in the uniformity of fluorescence across the surface of the membrane.

### Calculation of Retromer copies per cluster

#### Calibration of single fluorophore intensities

Fluorescently labeled proteins bound to a SLB were imaged using TIRF microscopy using continuous stream acquisition until nearly all spots bleached. Imaging started in a virgin region that was never exposed to excitation light. This was achieved by first focusing on a different location, turning on auto-focus, and moving to a new spot in dark. Thus, the fluorescent spots in first frame of a movie represent Retromer clusters that have not yet bleached. Laser power was adjusted to achieve stepwise bleaching in the illuminated sample. Each movie was corrected for uneven illumination field and background in a two-step process. First, the final frame in the movie, where almost all the fluorescent particles have bleached, was Gaussian blurred with a width of 2 pixels. Then this frame was subtracted from the entire image stack. Individual particles of fluorescence were identified and tracked using the ImageJ plugin SpeckleTrackerJ (37-39). Particles were automatically detected and tracked. Tracking typically started after some spots bleached; a lower spot density facilitated tracking. Tracks stopped the frame before a particle disappeared. This was checked visually, and tracks corrected if necessary. The tracks were saved and further analyzed in MATLAB. The last five frames before bleaching and five subsequent frames at the last position were analyzed by calculating the average pixel intensity in a 3×3 pixel region centered around the tracked position of the particle. The drop in intensity, *i*, due to this last bleaching step was calculated by subtracting the average of the 5 post-blech intensities from the 5 pre-bleach intensities. The distribution of *i* was plotted and checked that it fitted well with a Gaussian function. The mean single-fluorophore intensity, ⟨*i*⟩, was used to estimate the copy numbers of Retromer molecules in clusters detected in the first frame, as explained below. Each movie yielded 20-80 single-fluorophore intensity estimates from such bleaching events, and multiple image stack were analyzed for each individual experiment.

#### Analysis

We returned to the first frame in every movie and detected particles using SpeckleTrackerJ, employing the same criteria across different conditions. Using MATLAB, we calculated the mean pixel intensity for every spot, I, in a 3×3 pixel region around the spot’s centroid. Every spot’s intensity was divided by the average single-fluorophore intensity for that condition, ⟨*i*⟩, and the labeling efficiency, *e*, to estimate the number *N* of Retromer in that spot (cluster): *N = I/*(⟨*i*⟩ · *e*). Probability density functions of *N* are displayed with a bin width of 1. Typically, 3 movies per condition were tracked and analyzed for single-particle fluorescence, and 5 movies per condition were used for analysis of Retromer cluster size distributions. Two independent protein preparations were used per condition.

Statistical differences between distributions under separate conditions were tested using the two-sample Kolmogorov-Smirnov test (for comparing the distributions in Fig. 2. C and D) and the Kruskal-Wallis or one-way analysis of variance, followed by the multi-comparison tests (using Tukey’s Honestly Significant Difference Procedure, for comparing distributions in Fig. 4B or Fig. 5). These tests yielded low *p*-values in some cases, indicating the distributions are statistically different. Some differences are indeed obvious by inspection (e.g. Fig. 5 1:0 vs 50:1), but these differences lie mainly in changes in the monomer-to-oligomer ratios. The mean Retromer copies per cluster, and the spans varied within a limited range, 2.5 to 3.1 for the mean and [0-12] to [0-14] for the span. Thus, we do not find any evidence of biologically meaningful, large shifts in Retromer copy numbers across the conditions tested.

#### WASHC2C titrations

Five wells with identical conditions were set up with the specified protein components. WASHC2C was added to the well at the specified concentration and incubated for 30 minutes. The well was then washed a minimum of 3 times with Buffer A with 0.1% casein until bright spots of labeled WASHC2C were visible in the TIRF plane of the bilayer. Multiple bleaching movies were collected as described above and changes to the Retromer particle distribution were quantified. After analysis of the puncta of fluorescence in the micrographs (Fig. 4A), results were plotted as histograms (Fig. 4.A, B), with the order of the oligomer (i.e. 1 (monomers), 2 (dimers), 3 (tetramers), etc.) on the horizontal axis and relative frequency of the oligomer on the vertical axis. Statistical differences between the distributions were tested as described above.

## List of Abbreviations

AF488: AlexaFluor488®
AF546: AlexaFluor546®
AF647: AlexaFluor647®
DMT: Divalent Metal Transporter1-II (cargo)
WASHC2C-21: Fragment of WASHC2C with all 21 LFa motifs (aa357 to 1318)
WASHC2C-5: Fragment of WASHC2C with the final five LFa motifs (aa921 to 1318)
FRAP: Fluorescence Recovery After Photobleaching
LFa: Leucine-Phenylalanine-acidic residue motif
NiNTA-DGS: nickel salt of 1,2-dioleoyl-sn-glycero-3-[(N-(5-amino-1-carboxypentyl) iminodiacetic acid) succinyl]
RAB7-GTP: Rab7-Q67L, a constitutively GTP-bound form of Rab7
Retromer-RRS: Retromer with attached Retromer Recognition Sequence (RRS) of DMT cargo
RhPE: Rhodamine-phosphatidylethanolamine
SLB: Supported lipid bilayer
SNX3: Sorting nexin 3
SNX-BAR: Sorting nexin-Bin/Amphiphysin/Rvs
TIRF: Total Internal Reflection Fluorescence
VPS35: Vacuolar Protein Sorting-associated protein 35
VPS26: Vacuolar Protein Sorting-associated protein 26
VPS29: Vacuolar Protein Sorting-associated protein 29

## Acknowledgements

Research reported in this publication was supported by the National Institute of General Medical Sciences and the National Institute of Neurological Disorders and Stroke of the National Institutes of Health under awards GM060221 to CB, 1F32 GM125120-01A1 to CD, and R01NS113236 to EK.

## Figure Legends

**Figure S1.**
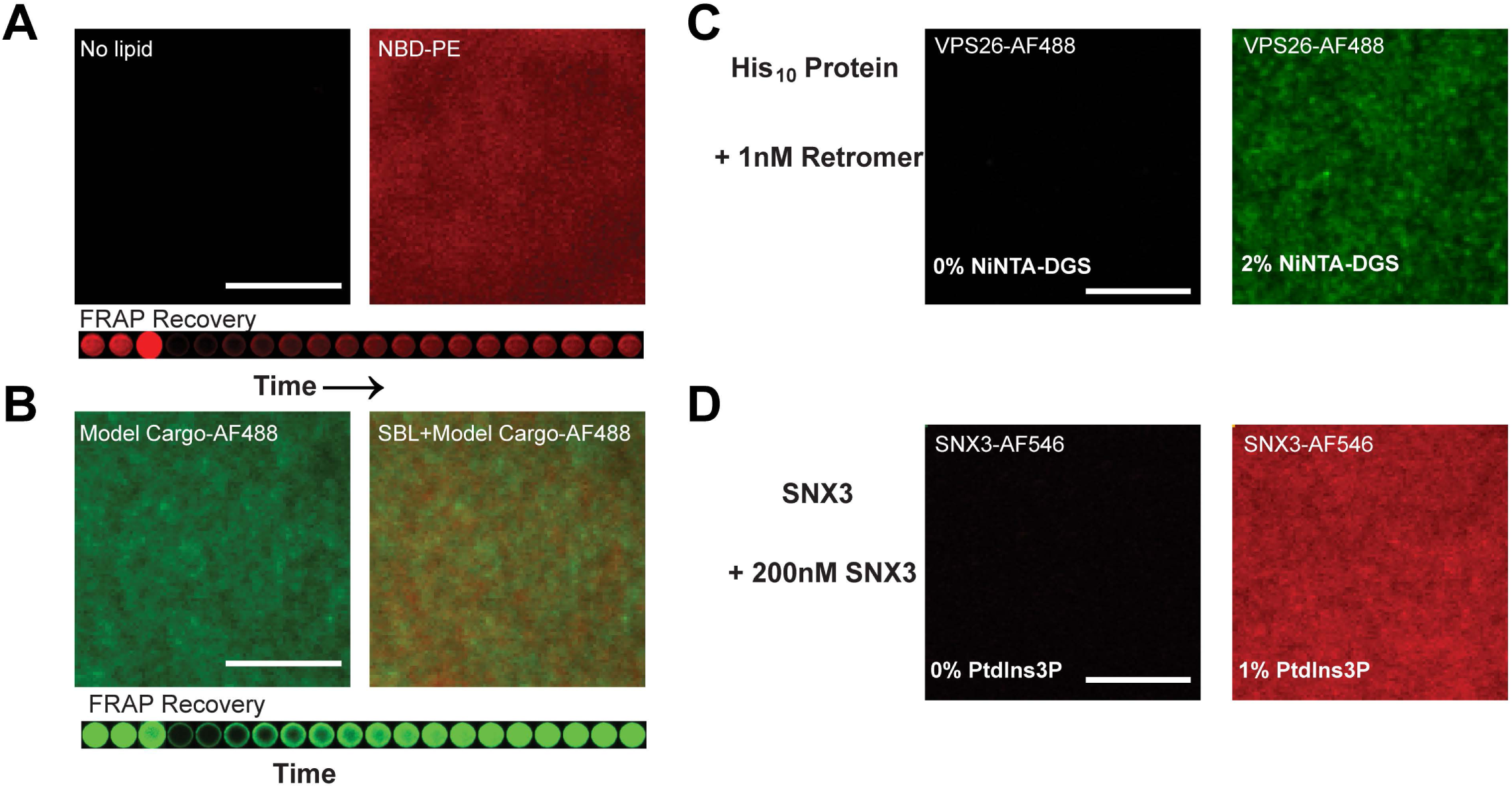
Quality of the SLB and specificity of protein attachment. **(A)**The SLB labeled with NBD-PE has a diffuse fluorescent signal (top), and recovers after FRAP (bottom), indicating that lipids in the bilayer are mobile. A lipid diffusion coefficient of 2.99 μm^2^/s is estimated from the recovery profile (36) **(B)** His-tagged, AF488-labeled model cargo protein was attached to the SLB (solution concentration 40 nM); attachment is dependent on the nickel lipid NiNTA-DGS. Cargo fluorescence is diffusely distributed on the surface of the SLB; bright puncta of fluorescence are absent (top panels). FRAP (bottom) shows that the cargo is mobile, with a diffusion coefficient of 0.88 μm^2^/s. **(C)** His-tagged, AF488-labeled Retromer attached to the SLB (solution concentration 1nM); attachment is dependent on nickel lipids. **(D)** AF546-labeled SNX3 attached to the SLB; attachment is PtdIns3*P* -dependent (solution concentration 200nm). Scale bars 10μm.

**Figure S2.**
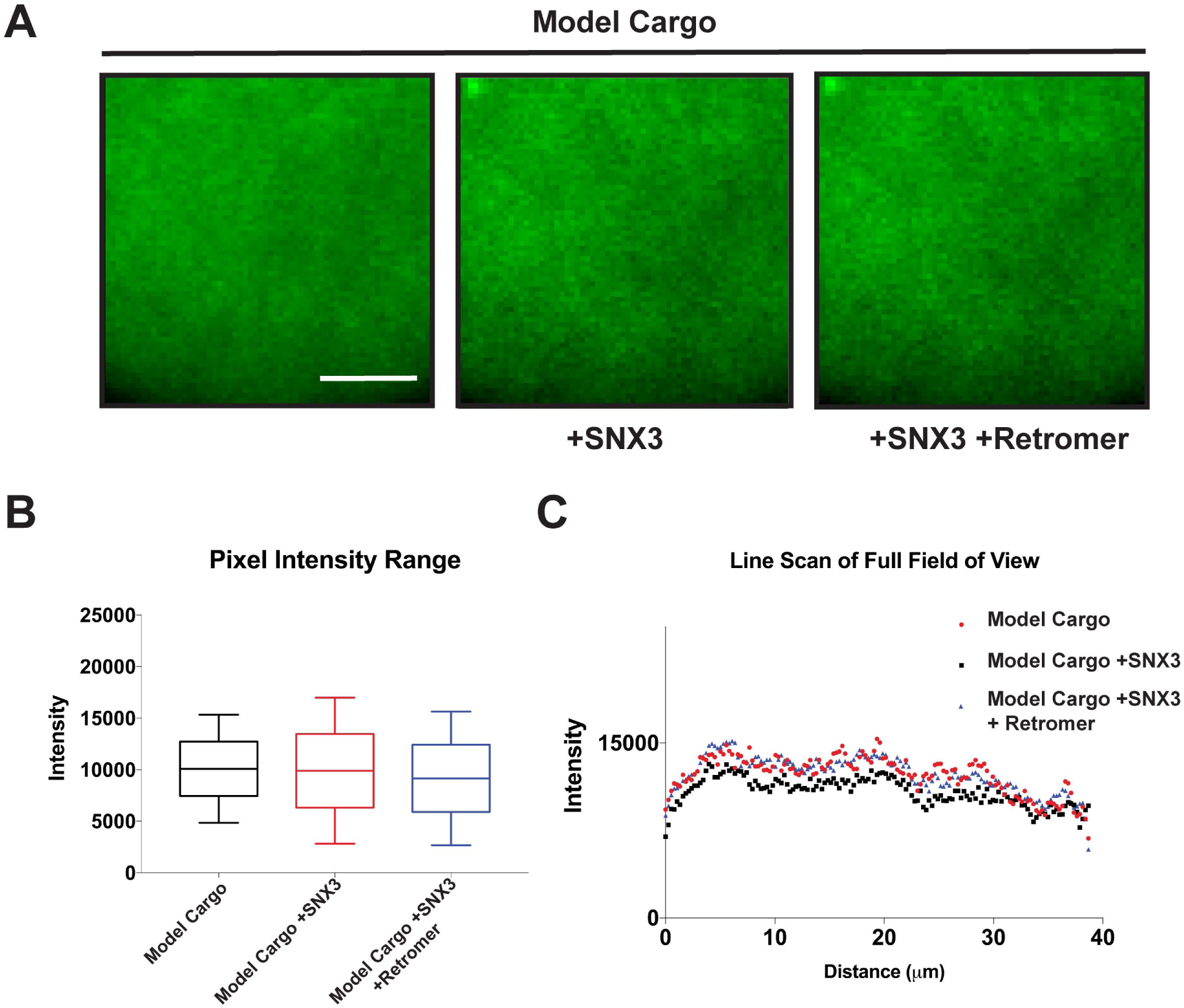
Retromer, SNX3, and cargo on the SLB. His-tagged, AF488-labeled model cargo protein was attached to the SLB at high protein density (solution concentration of cargo 40 nM). A.1) Model cargo protein alone, A.2) cargo and SNX3 (3μM), A.3) cargo, SNX3, and soluble (i.e. not his-tagged) Retromer (100 nM). All three conditions exhibit diffuse fluorescence across the SLB without bright puncta (B and C). Scale bar 5μm.

